# Thioflavin T indicates membrane potential in mammalian cells and can affect it in a blue light dependent manner

**DOI:** 10.1101/2021.10.22.465407

**Authors:** Emily Skates, Hadrien Delattre, Zoe Schofield, Munehiro Asally, Orkun S. Soyer

## Abstract

The fluorescent benzothiazole Thioflavin T (ThT) has a high binding affinity to protein aggregates and is used as a marker for the study of this process, most commonly in the context of neurodegenerative disease research and diagnosis. Recently, this same dye was shown to indicate membrane potential in bacteria due to its cationic nature. This finding prompted a question whether ThT fluorescence is linked to the membrane potential in mammalian cells, which would be important for appropriate utilisation of ThT in research and diagnosis. Here, we show that ThT localises into the mitochondria of HeLa cells in a membrane-potential dependent manner. Specifically, ThT colocalised in cells with a well-established mitochondrial membrane-potential indicator Tetramethylrhodamine methyl ester (TMRM) and gave similar temporal responses as TMRM to treatment with a protonophore, carbonyl cyanide-4-(trifluoromethoxy) phenylhydrazone (FCCP). Additionally, we found that presence of ThT together with exposure to blue light (λ=405 nm) exposure, but neither factor alone, caused depolarisation of mitochondrial membrane potential. This depolarisation effect was recapitulated by a mathematical model implementing the potential-dependent distribution of ThT and its light-dependent binding in mitochondria. These results show that ThT can act as a membrane potential dye in mammalian cells, when used at low concentrations and with low blue-light exposure, while it causes dissipation of the mitochondrial membrane potential at higher concentrations and in the presence of blue light excitation. This conclusion motivates a re-evaluation of ThT’s use at micromolar range in live-cell analyses, while indicating that this dye can enable future studies on the potential connections between membrane potential dynamics and protein aggregation.

## INTRODUCTION

Thioflavin T (ThT) is a fluorescent cationic benzothiazole dye that is widely used for quantification of amyloid fibril aggregation (Gade Malmos et al., 2017). These aggregates are shown to associate with a wide range of diseases such as Alzheimer’s, Parkinson’s, Type II diabetes and other age-related degenerative diseases (Xing, 2002; Xue et al., 2017), resulting in the wide use of ThT as a marker and research tool for studying these diseases (Groenning et al., 2007; Lockhart et al., 2005; Maezawa et al., 2008). The structure of ThT consists of a dimethylated benzothiazole ring coupled to a dimethylamino benzyl ring. In solution, these two rings act as a molecular rotor and their rotation around each other causes the low fluorescence emission of free ThT (Voropai et al., 2003; Stsiapura et al., 2007). Amyloid fibrils offer ThT a binding site, immobilizing its rotation, and thereby causing a characteristic increase in its fluorescence (Biancalana et al., 2009). ThT, by the same mechanism, can also exhibit increased fluorescence by binding to DNA (Biancardi et al., 2014), and RNA (Xu et al., 2016) and by forming micelles (Khurana et al., 2005).

Within the bacterial research community, it was shown that ThT can act as a Nernstian membrane potential indicator (Prindle et al., 2015; Stratford, et al., 2019) and can also influence bacterial membrane potential under certain conditions (Mancini et al., 2019; Sirec et al., 2019). In *Bacillus subtilis*, ThT distribution across the cell mimics that of 3-3’-dipropylthiadicarbocyanine iodide (DiSC_3_(5)), an established reporter for bacterial membrane potential (Geissler et al., 2000; Prindle et al., 2015), and of Tetramethylrhodamine, methyl ester (TMRM), a mammalian mitochondrial membrane potential dye that has also been used with bacteria (Sirec et al., 2019). These findings prompt a question whether the intracellular ThT distribution in mammalian cells may also follow mitochondrial membrane potential (ΔΨm), which would be important for appropriate utilisation of ThT in mammalian cell research and diagnosis.

Historically, the Nernstian equilibrium distribution of cationic lipophilic dyes, such as tetraphenylphosphonium (TPP) (Kamo et al., 1979) and the rhodamine dyes; e.g. rhodamine 123, TMRE and TMRM (Ehrenberg et al., 1988; Scaduto and Grotyohann, 1999), have been used as fluorescent markers for ΔΨm in mammalian cells. While plasma and mitochondrial potential are believed to be the main driver of the cellular distribution of these dyes, both dye distribution and fluorescence can also be altered upon binding to cellular components (Rottenberg, 1984; Ehrenberg et al., 1988). In turn, binding of the dyes to mitochondrial components is shown to be capable of reducing respiration activity, possibly due to membrane depolarisation (Scaduto and Grotyohann, 1999). Among the different cationic dyes studied, TMRM is most widely used because of its low binding to cellular components, a high fluorescence signal, and rapid and reversible equilibration across the membranes (Scaduto and Grotyohann, 1999). Given its cationic nature, ThT would also be expected to distribute itself in cells according to the Nernst equation, but possibly with its distribution also influenced by its ability to bind to macromolecules such as protein aggregates, DNA, and RNA.

Here, we analysed ThT dynamics in HeLa cells as a model mammalian system. We found that ThT, when applied at low micromolar concentrations and with low blue-light exposure, distributes in the cell according to ΔΨm. In particular, ThT co-localized in cells with TMRM and gave similar temporal responses as TMRM to perturbance of ΔΨm. With increased concentrations, and when cells were under blue light (λ=405 nm) exposure, ThT also caused a depolarisation of the mitochondrial membrane. We show that these findings can be explained by a simple mathematical model that incorporates potential-dependent distribution of ThT and assumes a light-dependent binding behaviour. Taken together, these results show that ThT can act as a ΔΨm indicator dye in mammalian cells but can also dissipate ΔΨm in a manner dependent on both concentration and blue light. While the latter finding cautions for use of ThT at high concentrations and under light exposure, the former finding opens the possibility for utilising ThT for studying ΔΨm dynamics especially in microfluidic devices, where TMRM and other membrane-potential indicators have undesired high affinity to PDMS which gives rise to high background fluorescence (Zand et al., 2013) while ThT doesn’t (Prindle et al. 2015).

## RESULTS

### ThT distributes in mitochondria at low concentrations

ThT is most commonly used at ∼10 μM for the analysis of protein aggregates (Xue et al., 2017). To evaluate the membrane-potential dependency of ThT distribution at the single-cell level, we incubated HeLa cells in media supplemented with 0.2, 0.5 and 1 μM ThT for an hour at 37°C and imaged them by scanning confocal microscopy using a 405-nm laser for excitation (Fig. 1A and B). The results suggested that not only the fluorescence level but also the spatial distribution of ThT varies with concentration. At 1 μM, ThT fluorescence was shown throughout the cell and particularly localised in the nucleoli. At 0.2 and 0.5 μM, in contrast, ThT localised in a pattern reminiscent of the mitochondrial network. To further examine if ThT localises in mitochondria when used at low concentrations, we co-stained HeLa cells with the mitochondrial membrane-potential indicator TMRM and found that the spatial distribution of ThT overlapped with that of TMRM (Fig. 1C and D). This co-localisation with TMRM, and the aforementioned cationic nature of ThT, suggests that ThT distributes itself according to the electrical potential differences within the cell and therefore might respond to changes in the ΔΨm.

**Figure 1:**
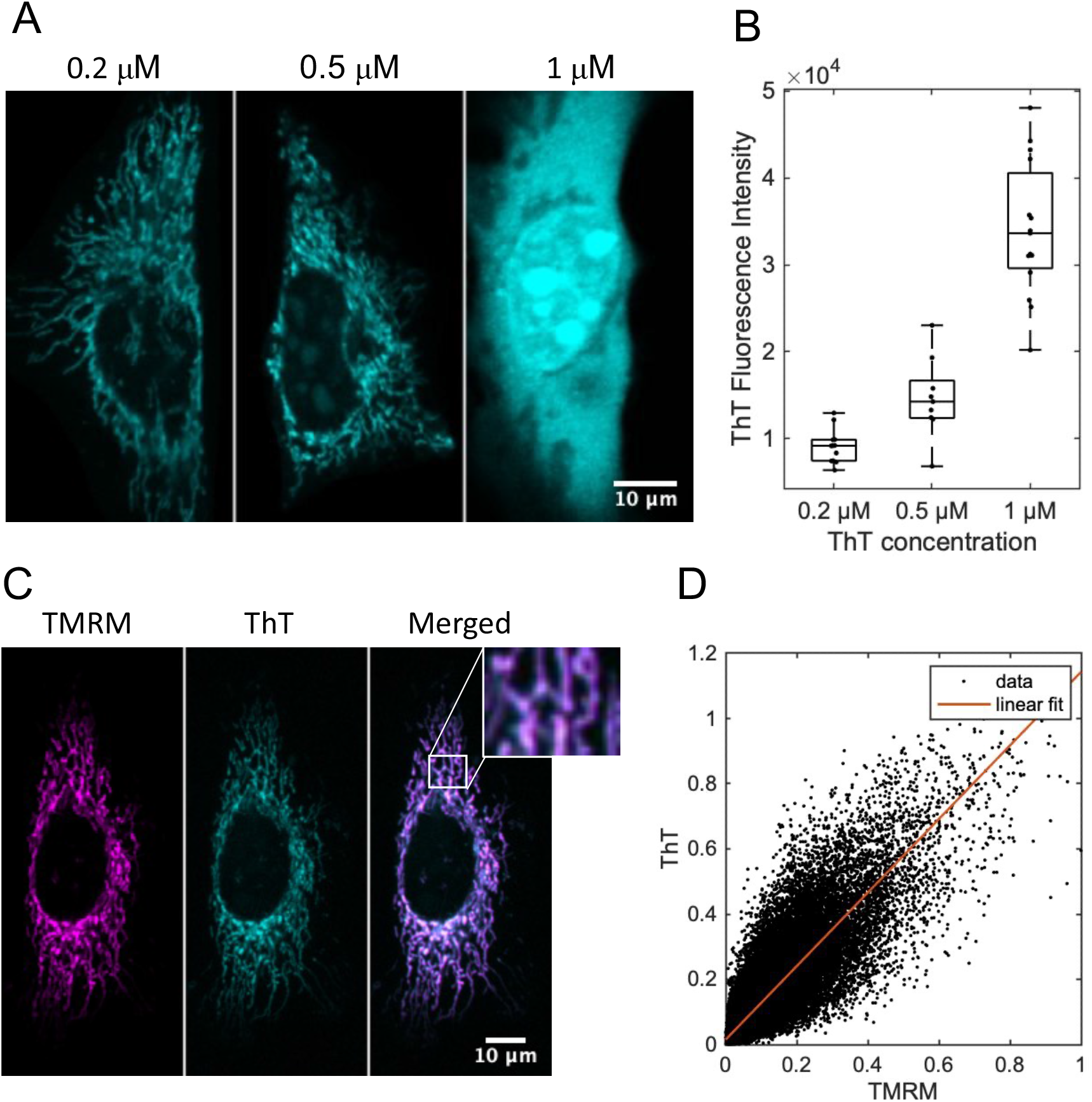
**(A)** Distribution of ThT in HeLa cells at ThT concentrations of 0.2, 0.5, and 1 μM (left to right). The scale bar applies to all images. Excitation was at 405 nm (see *Methods* for further details of imaging conditions). Each cell is representative of the total population. **(B)** Fluorescence intensity of a population of cells at ThT concentrations of 0.2, 0.5 and 1 μM. A total of 12, 9 and 15 cells were analysed at the respective concentrations. Differences among populations were statistically significant (U test, p < 0.05). **(C)** Separate and overlayed images of a ThT and TMRM stained HeLa cell. The dye concentrations used were 0.2 μM and 25 nM, for ThT and TMRM, respectively. The insert shows an enlarged section of the mitochondrial network. The colour scheme, as a function of pixel intensity, is applied afterwards as an aid to the eye. The scale bar applies to all images. **(D)** Correlation of pixel-wise fluorescence intensity from TMRM and ThT images, and a corresponding fit using a linear model shown as a red line (R^2^ = 0.8858).

### ThT responds to changes in mitochondrial membrane potential

To test the membrane-potential dependency of ThT localisation in the mitochondria, we recorded ThT distribution under chemical perturbation of ΔΨm by FCCP, a protonophore that facilitates the passive transport of protons across the inner mitochondrial membrane, and thereby causing a collapse of ΔΨm. It has been shown that depolarisation by FCCP results in the loss of cationic dye fluorescence from the mitochondria within 5-10 minutes (Farkas et al., 1989; Nicholls, 2012). Following this approach, we performed fluorescence time-lapse microscopy of HeLa cells for 9 minutes, during which 2 μM FCCP was added to the media at 2 minutes into the experiment. We then quantified the fluorescence intensity of TMRM or ThT over time, for individual cells, and plotted the rate of change from pre-FCCP treatment.

Consistent with a previous study (Nicholls, 2012), addition of FCCP caused a reduction in TMRM fluorescence, while mock control did not show any significant changes (Fig. 2A-C). Having confirmed the dynamics with TMRM in our experimental setup and with HeLa cells, we repeated this experiment with ThT. The fluorescence intensity of ThT displayed similar dynamics as with TMRM, with fluorescence showing a decrease after the addition of FCCP (Fig. 2D-F). We note that there is less of a decrease in ThT fluorescence with FCCP addition, compared to TMRM. This is partially because, after FCCP addition, ThT distributes in the cytoplasm and localizes in the nucleoli, while TMRM had mostly leaves the cells. These results suggest that while the mitochondrial localisation of ThT is dependent on the ΔΨm, there are additional effects likely due to ThT binding in the cell.

**Figure 2:**
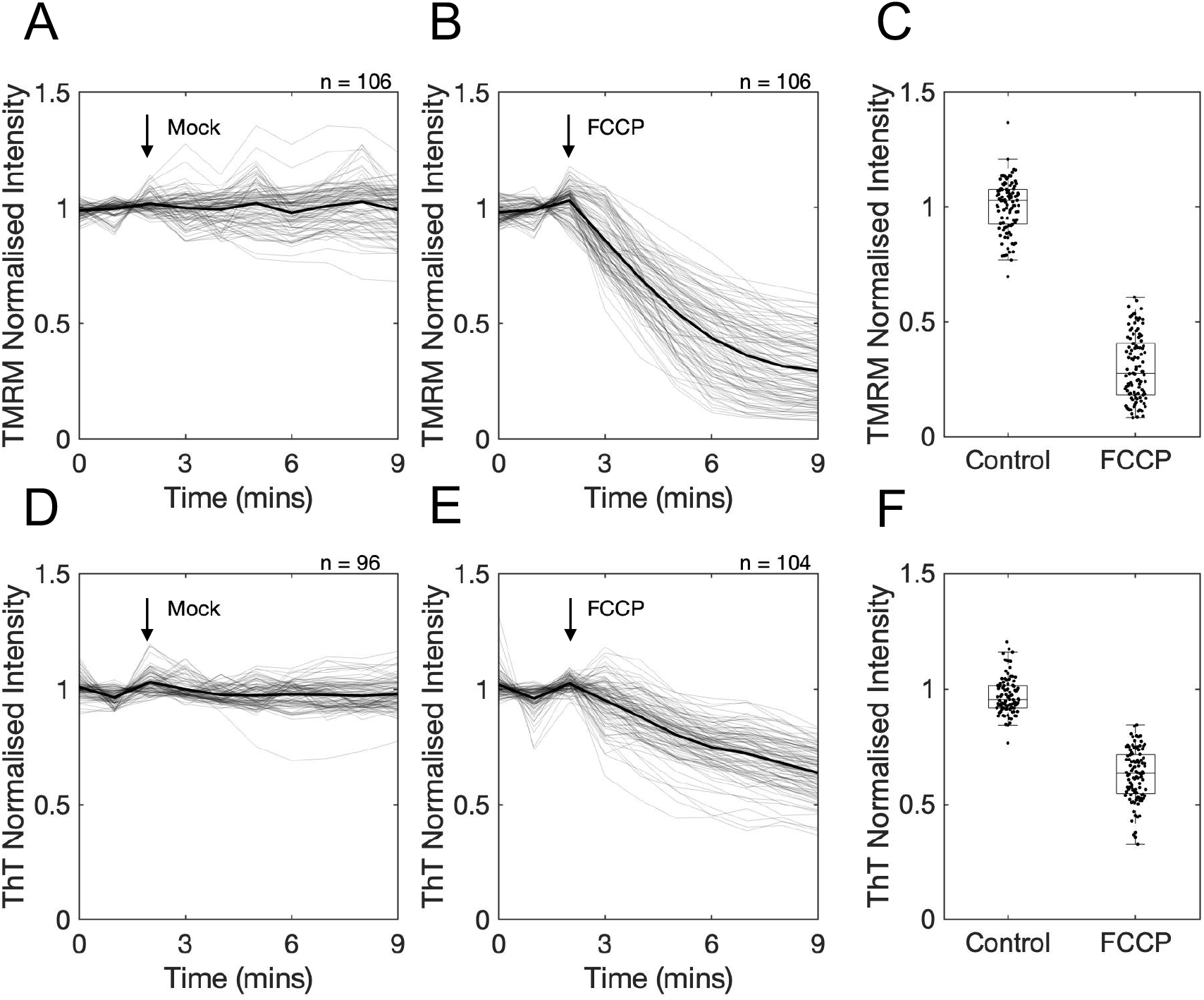
**(A-B)** The temporal response of cellular TMRM fluorescence, after the addition of cell culture media as a control **(A)** or media containing 2 μM FCCP **(B)**. Cells were incubated with 25nM TMRM and the TMRM fluorescence was normalised to that from the first image. The point of media addition is indicated with an arrow. On each panel, the grey lines show the fluorescence intensity from single cells, and the black line shows the population mean. Data is collated from three independent experiments, each with three technical repeats, resulting in a total of 106 cells analysed in both A and B. **(C)** The comparison of the final fluorescence intensity (time point 9mins) shown on panels A and B (i.e. without and with FCCP addition). Data from the entire cell population is shown as box-plots. The means of the two distributions are statistically significant (U test, p < 0.05). **(D-E)** The response of cellular ThT fluorescence, after the addition of cell culture media as a control **(D)** or media containing 2 μM FCCP **(E)**. Cells were incubated with 0.2 μM ThT and the ThT fluorescence was normalised to that from the first image. Experiment design is same as for TMRM, resulting in a total of 96 and 104 cells analysed in D and E, respectively. **(F)** The comparison of the data shown on panels D and E, done in the same way as explained in C. The means of the two distributions are statistically significant (U test, p < 0.05).

### ThT affects mitochondrial membrane potential in a concentration and blue-light dependent manner

While the above results show that ThT can distribute in mitochondria in a ΔΨm dependent manner, we also observe ThT distributing throughout the cell and in nucleoli when used at 1 μM (Fig. 1A). These two results could be corroborated under the hypothesis that ThT affects the ΔΨm at high concentration, leading to ΔΨm dissipation and ThT leaving mitochondria. Indeed, previous studies have shown that the binding of membrane-potential dyes to mitochondrial components could dissipate ΔΨm (Scaduto & Grotyohann, 1999). To test this possibility for ThT, we cultured cells with media containing different concentrations of ThT while monitoring ΔΨm with TMRM (Fig. 3A). We used TMRM at a low concentration (25 nM) to ensure that TMRM itself does not influence the ΔΨm (Scaduto & Grotyohann, 1999). Cells were incubated with the dyes for 1 hour at 37°C and then imaged by fluorescence microscopy every 15 secs for 20 mins. At a ThT concentration of 0.2 μM, we saw no change in ΔΨm, as monitored by TMRM fluorescence (Fig. 3B, dashed line. See also Fig. S1). However, at ThT concentrations of 1 and 5 μM, a decrease in TMRM fluorescence was observed (Fig. 3B, dash-dot and dotted lines. See also Fig. S1). This suggests that there is a concentration-dependent loss of ΔΨm due to ThT. In line with this view, we also found that at 1 and 5 μM, ThT leaves the mitochondria and localises in the nucleoli (Fig. S2A). The time it takes for the mitochondrial membrane to depolarise was shorter at higher ThT concentrations (Fig. 3B).

**Figure 3:**
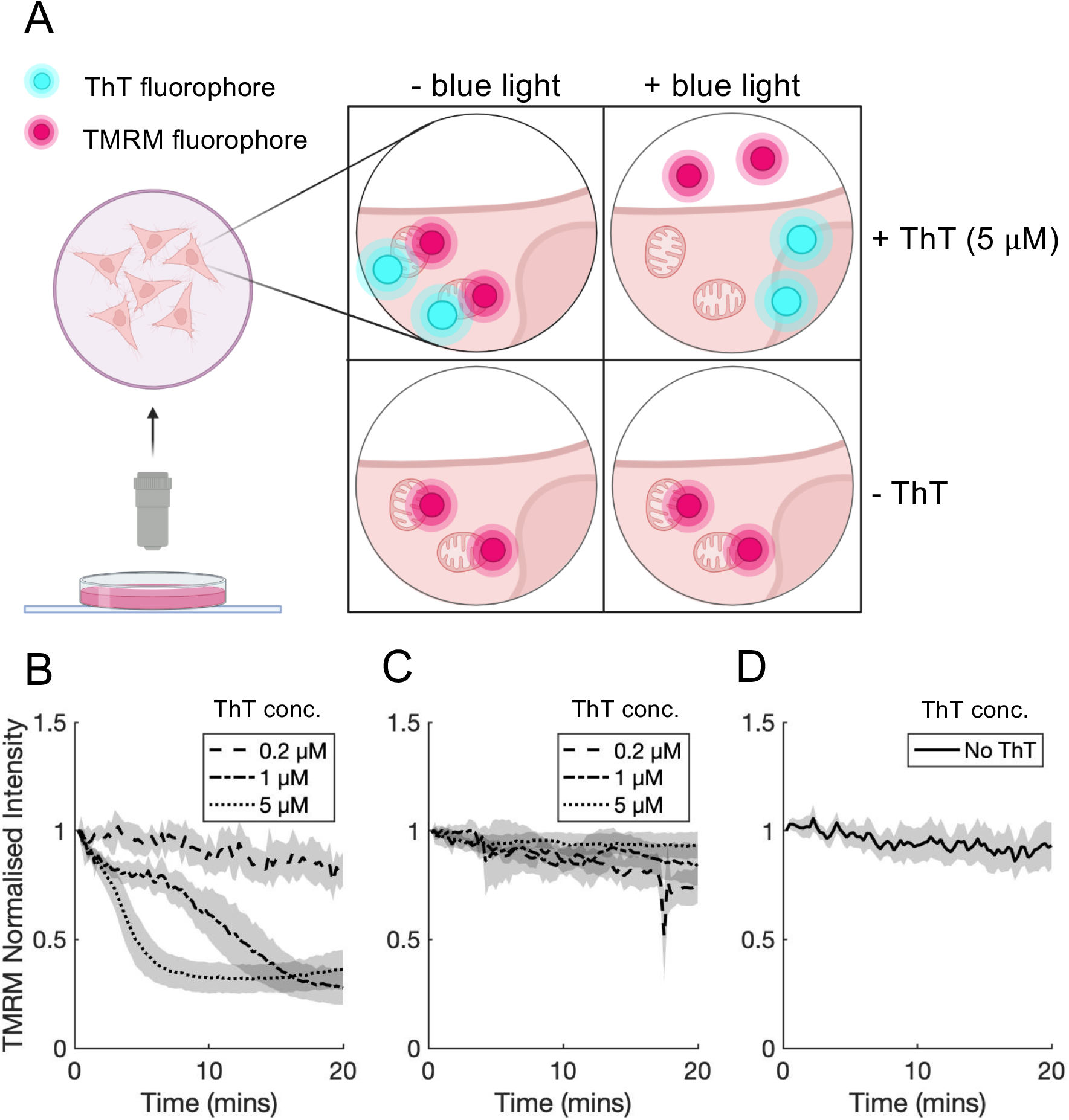
**(A)** Cartoon depicting the setup for the co-staining experiments. Cells were incubated with either TMRM only or with TMRM and ThT, and then imaged with either red light only or with red and blue light. Cartoon created with BioRender.com. **(B-C)** Temporal TMRM fluorescence from cells incubated with TMRM (25 nM) and co-incubated with 0.2, 1 and 5 μM ThT, imaged in either red and blue light **(B)** or just red light **(C**). On each panel, the lines show the mean fluorescence intensity with the shaded section showing population standard deviation, SD (mean±SD). Data is collated from three independent experiments, each with three technical repeats resulting in a total of 176 and 177 cells analysed in B and C. **(D)** Temporal TMRM fluorescence from cells incubated with TMRM (25 nM) without ThT co-staining and imaged in red and blue light. Experiment design is same as for B-C, resulting in a total of 62 cells.

While these results suggest that higher ThT concentrations cause a loss of ΔΨm, it is also possible that this effect arises from the blue light exposure that is used for imaging ThT. Indeed, it has been shown that blue light can cause a multitude of effects on cell integrity and physiology (Denning et al., 2002; Ramakrishnan et al., 2016; Waldchen et al., 2015). Preliminary results indicated a possible role for blue light, as we had found no loss in TMRM fluorescence (i.e. no change in ΔΨm), when cells were imaged at the beginning of a time-lapse experiment and after a 1-hour pre-incubation with both dyes, regardless of the ThT concentration (Fig. S3). To distinguish between the possibilities of either ThT and/or blue light causing the observed loss of ΔΨm, we repeated the co-incubation, time-lapse experiment, but imaged the cells with the 561-nm laser only for 20 mins (i.e. no blue light exposure – see Fig. 3A). We took ThT images only at the beginning and end of this time-lapse experiment, to assess the ThT distribution. Over the course of the time lapse, TMRM fluorescence was stable (Fig. 3C), and no nucleoli localisation of ThT was observed at the end of the experiment (Fig. S4). This suggests that the presence of ThT in the cell on its own does not cause a loss of ΔΨm. We then tested whether blue light exposure during our imaging is sufficient on its own to cause membrane depolarisation. Cells were incubated with TMRM only and were imaged using both red and blue light every 15 secs for 20 min, as above. We again found no effect on TMRM fluorescence and therefore on ΔΨm (Fig. 3D). These results strongly suggest that a combination of ThT and blue-light exposure is needed to cause a loss of ΔΨm, which then leads to both TMRM and ThT exit from mitochondria, and subsequent localisation of the latter into nucleoli (Fig. S1A).

### ThT and blue light effects are explained by a simple mathematical model

To explain and better understand these experimental results, we aimed to develop a simple mechanistic model. The aim of the model was to see what mechanistic assumptions need to be made to reproduce the key experimental observation of the combined need for blue light and ThT to cause depolarisation of the membrane potential. We created a two-compartment cell model to represent inside and outside of the mitochondria (see *Methods* and *Supplementary File* 1 and 2 for a MATLAB implementation of the model). The model implements passive flux of ThT and TMRM across both plasma and mitochondrial membranes according to the compartment potentials and dye concentration, using the Goldman-Hodgkin-Katz (GHK) flux equation (Goldman, 1943; Hodgkin & Katz, 1949), and dynamic binding of ThT in each compartment (Fig. 4A).

**Figure 4:**
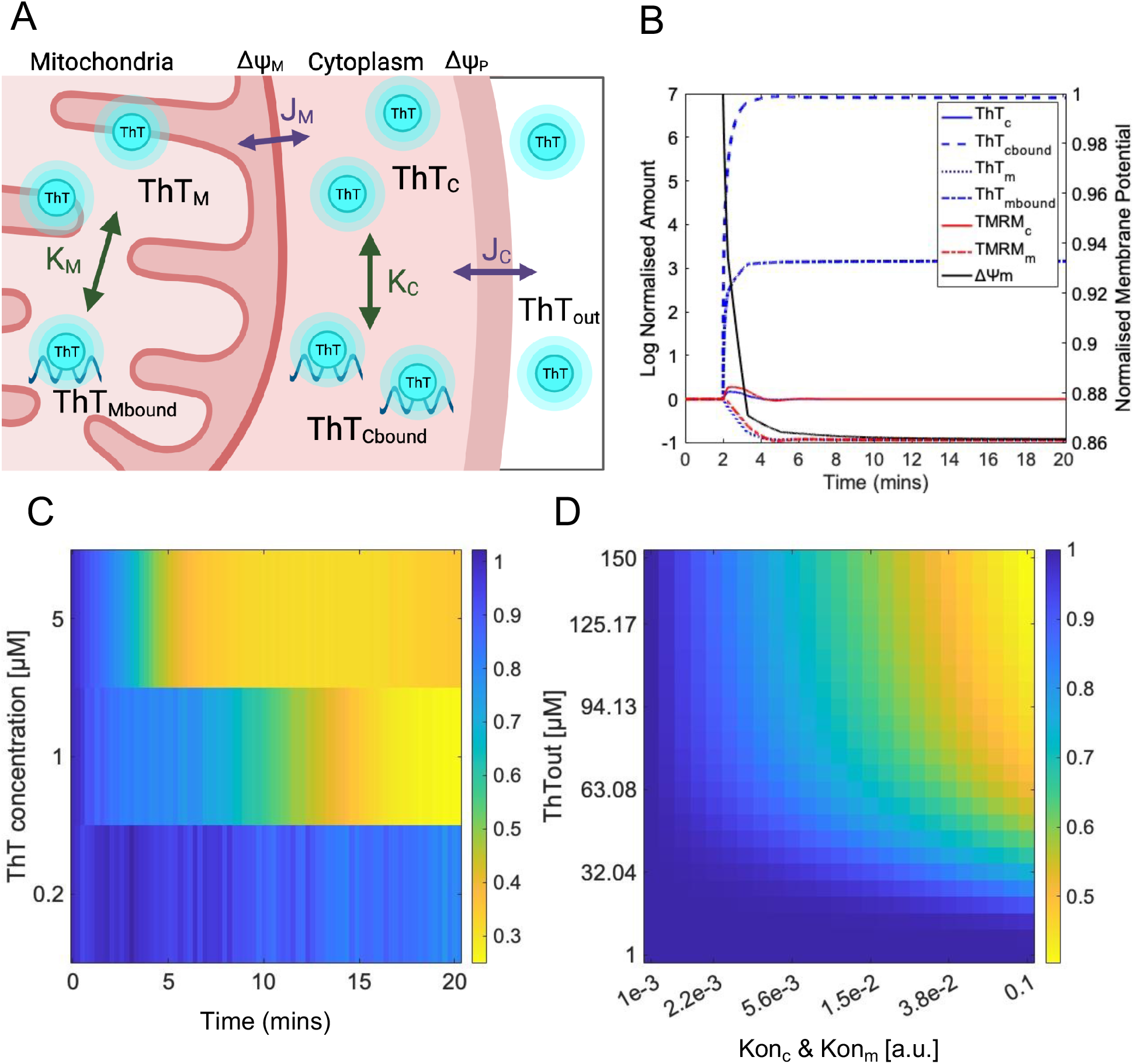
**(A)** Cartoon depicting the mathematical model, which features two compartments as mitochondria and cytosol. The different simulated variables are shown as described in the main text. Arrows indicate binding events and dye flux across membranes. Cartoon created with BioRender.com. **(B)** Normalised concentrations of simulated variables of the model over time. Simulation is run to steady state and then the system is perturbed at t=2mins by increasing the parameters *Kon*_*m*_ and *Kon*_*c*_ by 1000-fold. **(C)** Phase plot showing the fold-change in TMRM fluorescence (colour coded) over time (x-axis), during time-lapse experiments with different levels of ThT (y-axis). This data is the same as shown in Fig 3B but is compiled here into a phase plot to allow easier comparison to simulations. **(D)** Phase plot showing normalised steady-state TMRM level (colour coded) from simulations performed with different ThT binding coefficient (x-axis, mimicking increasing light exposure) and different levels of ThT (y-axis).

We found that to reproduce the combined effects of ThT and the light, the model needed to implement a dependence of ΔΨm on the concentration of the bound form of ThT in the mitochondria (*ThT*_*mbound*_) and a light-dependence of ThT binding dynamics (see *Methods*). The former assumption is supported by the dissipation of ΔΨm by the binding of cationic dyes in the mitochondria when used at higher concentrations (Scaduto & Grotyohann, 1999), while the latter assumption reflects an expected increase in unfolded proteins or other binding sites for ThT upon light exposure (Redecke et al., 2009). Using this model, we simulated the distribution of both ThT and TMRM (Fig. 4B). In these simulations, dyes equilibrate across the two compartments in a membrane potential dependent manner, as expected by the GHK flux equation. When we simulate conditions that mimic absence of blue light (i.e. the binding coefficients for ThT in the mitochondria, *Kon*_*m*_ and cytosol, *Kon*_*c*_, are set to a low value), the dye equilibration does not impact ΔΨm significantly, which reaches a steady state close to its initial value set at the beginning of the simulation (Fig. 4B, t<2mins). The main reason for this is that at low ThT binding throughout the cell, there is little *ThT*_*mbound*_ and therefore almost no effect on ΔΨm nor on ΔΨm related steady-state levels of the dye. When we simulate blue light (i.e. setting a high value for *Kon*_*m*_ and *Kon*_*c*_), there is an increase in the amount of both *ThT*_*mbound*_ and *ThT*_*cbound*_ (Fig. 4B, simulation traces after t=2mins). These dynamic changes in ThT distribution cause a depolarization of ΔΨm, reaching a new steady state at a less negative potential, and consequently of the TMRM leaving the cell (Fig. 4B).

Our experimental results show that at low concentrations of ThT, blue light had no effect on TMRM fluorescence and therefore on the ΔΨm. To see if our model can recapture this result, we ran simulations with different concentrations of ThT and at different values of *Kon*_*m*_ and *Kon*_*c*_, mimicking different light exposure levels. We then plotted the resulting TMRM concentration from these simulations against ThT concentration and light exposure level used, creating a phase plot for the system dynamics (Fig 4C). This phase plot shows that TMRM concentration in mitochondria is dependent on both ThT level and light exposure, as seen experimentally. The simulation-based results of Fig. 4C are driven by the basal ThT levels in the mitochondria (determined by the overall ThT concentration, y-axis), the nonlinear binding dynamics of ThT in the mitochondria (determined by light exposure, x-axis) and the interplay between *ThT*_*mbound*_ concentration and ΔΨm. The phase plot demonstrates that increasing light will only cause a substantial effect on ΔΨm, when overall ThT concentration is above a certain level, and conversely, the higher the ThT concentration, less blue light exposure (increase in *Kon*_*m*_ and *Kon*_*c*_) is required for the depolarisation of ΔΨm. This is in line with our experimental results demonstrating that less blue light is required for depolarisation of the mitochondrial membrane at higher ThT concentrations. Altogether, these results show that a model incorporating key mechanistic assumptions about ThT binding in mitochondria and cytosol in a light dependent manner and a dependence of ΔΨm on *ThT*_*mbound*_ can qualitatively explain the observed experimental results.

## DISCUSSION

We investigated the possibility that ThT, a commonly used protein-aggregate marker, also acts as ΔΨm indicator in a mammalian cell. Using live single-cell microscopy we show that ThT can distribute itself in mitochondria in a ΔΨm-dependent manner and that it responds to mitochondrial membrane depolarisation by leaving mitochondria. Both responses show similar qualitative dynamics to a well-established ΔΨm indicator, TMRM. Unlike TMRM, however, ThT can also accumulate in the nucleoli upon mitochondrial membrane depolarisation, presumably due to high binding affinity of ThT to RNA and protein aggregates which are abundant in nucleolus (Latonen, 2019). Investigating this accumulation dynamics further, we found that ThT, in a concentration and blue light exposure dependent manner, can cause mitochondrial membrane depolarisation. Neither ThT nor blue light exposure alone had a similar effect. Based on these findings, we show that a simple mathematical model incorporating ΔΨm-based ThT distribution and light-dependent ThT binding in mitochondria could qualitatively recapitulate the experimental observations.

ThT has been used for live-cell time-lapse imaging of protein aggregates (e.g., Beriault & Werstuck, 2013; Caron et al., 2014; Sugimoto et al., 2015). Our finding, that the blue-light exposure for imaging ThT fluorescence could impact ΔΨm, raises a cautionary note for the use of ThT in live cell imaging for assessment of protein aggregates or DNA. Particularly, we note that changes in ΔΨm could alter cellular ATP levels, which is shown to interlink to the prevention of protein aggregation through ATP’s destabilising effects on aggregates (Patel et al., 2017; Pu et al., 2019). This suggests that changes in ΔΨm, caused by ThT presence and blue light imaging, can then feedback to alter protein aggregation levels. Therefore, the interpretations of live-cell ThT microscopy, regardless of whether it is aimed to study protein aggregates or ΔΨm, would need careful consideration of the possible effects arising from mitochondrial membrane potential depolarisation. We note that neither ThT, nor blue light on their own, was found to impact ΔΨm in our experiments, suggesting that ThT can still be utilised for live-cell studies when it is conducted with low concentration of ThT and with low exposure imaging modalities.

In our model, ThT binding in mitochondria was assumed to be light dependent. This assumption was motivated by the fact that blue light can cause mitochondrial DNA damage (Godley et al., 2005), which may increase the binding sites for ThT there. However, the exact biophysical and biochemical mechanism of this is unclear. Elucidating this mechanism in a future project utilising ThT as a unique dye would be important for understanding the link between bioenergetics and intracellular molecular aggregation. It may also provide a useful experimental tool to modulate the mitochondrial membrane potential in a spatially and temporally controlled manner using light.

The finding that ThT can act as a ΔΨm indicator opens up new directions of study. One particular area is the use of ThT to study ΔΨm and its dynamics in mammalian cells using microfluidics. Most microfluidic channels are made from polydimethylsiloxane (PDMS), which is shown to bind TMRM and thereby cause a high background signal. This limits the utilisation of TMRM in microfluidic studies, as it makes it very hard to distinguish between cells and background (Zand et al., 2013). ThT, has been previously used in microfluidics and doesn’t not present the same issues (Saar et al., 2016; Shammas et al., 2015; Prindle et al., 2015). Combined with the presented findings, this opens up a potential opportunity to take ΔΨm measurements of single mammalian cells in microfluidic devices.

## METHODS

### Cell culture

The HeLa cells were sourced from the Public Health England (ECACC catalogue no.93021013). The cells were kept as cryo-stocks and live cultures, where the latter was never passaged more than 10 times or for longer than 3 months as stated in the guidelines for cell culturing (Geraghty et al., 2014). Cultures were maintained in minimum essential media (MEM) with NaHCO_3_ (Sigma, M2276), supplemented with 1% L-glutamine (Sigma, G7513), 10% heat-inactivated fetal calf serum (HIFC), 1% non-essential amino acids (Sigma, M7145), and 1% penicillin/streptomycin solution (Sigma, P4333), and stored in a humidified atmosphere in 5% CO_2_ at 37°C. Cells for fluorescence microscopy were seeded at 2×10^5^ in glass bottom 6-well plates (MatTek, P06G-1.5-10 F) and cultured until ∼70% confluency. For microscopy, cells were first washed with PBS and then placed in MEM without NaHCO_3_, and buffered with 10% HEPES (Sigma, H0887) instead, and supplemented with 10% HIFC (ThermoFisher Scientific, 1008-2147) and 1% penicillin/streptomycin solution (Sigma, P4333).

### Dyes and chemical reagents

Thioflavin T (ThT, Sigma Aldrich T-3516) was kept as a 10 mM stock solution in distilled water and stored at 4°C in the dark. TMRM (ThermoFisher Scientific, T-668) was dissolved in DMSO to make a 10 mM stock, which was stored at −20°C in the dark. Carbonyl cyanide 4-(trifluoromethoxy)phenylhydrazone (FCCP, Abcam ab120081) was dissolved in DMSO to make a 1 mM stock and stored at −20°C in the dark.

### Fluorescence microscopy

All images and time-lapse videos were recorded using a laser scanning confocal microscope (LSM-880, Zeiss, Oberkochen, Germany) with an EC Plan-Neofluar 40x/1.30 Oil DIC M27 objective lens unless stated otherwise. After the addition of either fluorescence dyes, cells were incubated in HEPES buffered media at 37°C with ambient atmosphere for an hour in the dark to allow for the equilibration of the dye and cells. ThT and TMRM dyes were imaged using a 405- and 561-nm laser, respectively. The maximum power output for both lasers was 250 mW and they were applied at a 5% power setting. Light intensity was measured at the sample plane using a power meter (PM160T, Thorlabs, Inc., New Jersey) for each laser, giving 48.4 and 27.7 μW for the 561 and 405 nm laser, respectively.

For the assessment of the localisation of ThT dye at different concentrations, cells were incubated with 0.2, 1 and 5 μM of ThT for an hour prior to imaging. For the co-localisation assay, cells were incubated for an hour with 25 nM TMRM and 0.2 μM ThT prior to imaging. For these experiments, cells were imaged using a Plan-Apochromat 63x/1.40 Oil DIC M27 objective lens (Zeiss, Oberkochen, Germany) and frame averaging was set to 16 to improve signal to noise ratio (i.e. 16 images were taken and averaged over each pixel).

To assess both dyes’ response to changes in the mitochondrial membrane potential, FCCP at final concentration of 2 μM was used to artificially depolarise the mitochondrial membrane potential and the response of the dye was monitored by time-lapse microscopy. For the duration of the experiment the temperature was set to 37°C in the microscopy chamber. A 9-minute long time-lapse was set, so to take images every minute. Time-lapse was paused just before the 3^rd^ frame and 1 mL of media solution supplemented with dye and FCCP was added. The time-lapse was then restarted immediately. For the control experiments a mock injection was made with media and dye only.

### Image analysis

Image segmentation was achieved using Fiji/ImageJ (National Institutes of Health) (Schindelin et al., 2012) where the cytoplasmic region for each cell was manually selected and the mean fluorescence intensity values were obtained. Figures were created using MATLAB (MathWorks). Fluorescence data from time-lapse experiments was obtained as above and then normalised for each cell by the average mean fluorescence of the first three frames, before the addition of FCCP. The boxplots were generated by plotting the ratio between the fluorescence of the first frame and last frame, and the individual data points were plotted using the plot spread points (beeswarm plots) algorithm (Jonas, 2021). For colocalization analysis, both ThT and TMRM fluorescence data were normalised using a min/max normalisation. A scatter plot generated to which a linear model was fit, and the Pearson correlation coefficient calculated using the appropriate MATLAB functions.

### Mathematical model

The model presented here considered the cell as a two-compartment system separated by membranes. Thus, our abstract cell has a mitochondrial and a cytosolic compartment and is separated from the extracellular environment by a plasma membrane. The model accounted for potential-driven, passive diffusion of two cationic dyes across these compartments, using the Goldman-Hodgkin-Katz flux equation (Goldman, 1943; Hodgkin & Katz, 1949). For ThT, the model incorporated binding in both compartments. Finally, the model assumed a simple, linear relation between mitochondrial bound ThT level and ΔΨm. The model thus consisted of the following ordinary differential equations (ODEs) describing the temporal dynamics of mitochondrial and cytosolic TMRM (*TMRM*_*m*_ and *TMRM*_*c*_), and mitochondrial and cytosolic free and bound ThT (*ThT*_*m*_, *ThT*_*c*_, *ThT*_*c_bound*_, and *ThT*_*m_bound*_):

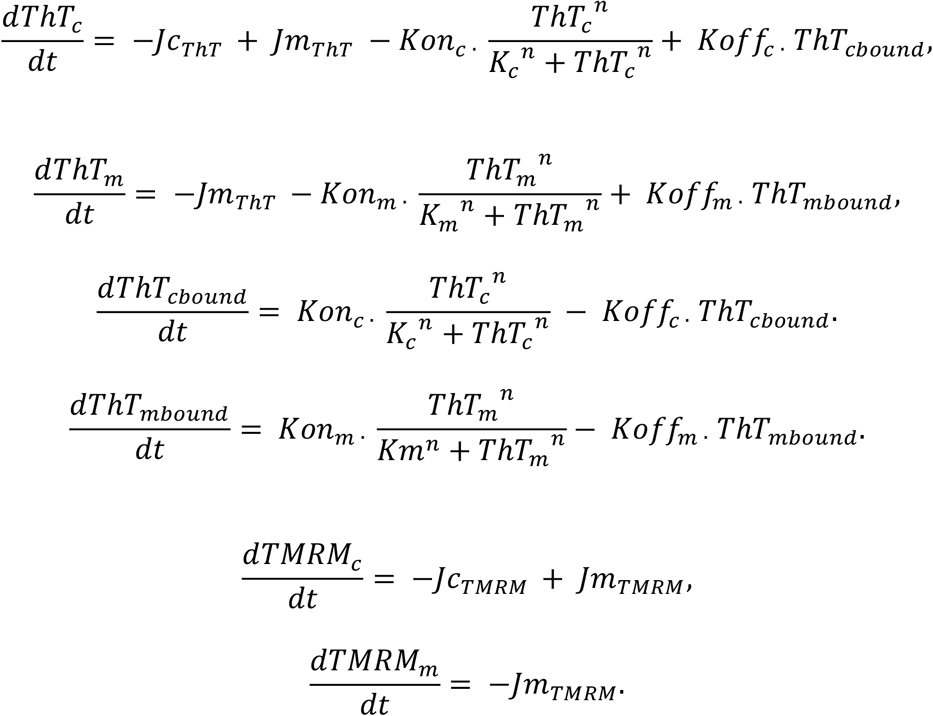

where the passive, potential-driven fluxes of ThT and TMRM across the plasma and mitochondria membranes are described by the terms denoted by *Jc*_*ThT*_ and *Jm*_*ThT*_ and by *JTMRM* and *Jm*_*TMRM*_ respectively. The binding of ThT in the cytoplasm and mitochondria are both modelled using a cooperative Hill-function where *n* is the cooperativity coefficient. *K*_*m*_ and *K*_*c*_ are the ThT concentrations producing half occupancy in cytosol and mitochondria respectively, and *Kon*_*c*_ and *Kon*_*m*_ are the maximal rate of binding in these two compartments. The dissociation of cytoplasmic and mitochondrial bound ThT is determined by the dissociation constants, *Koff*_*c*_ and *Koff*_*m*_ respectively.

The generic form of the Goldman-Hodgkin-Katz flux equation is used to determine the *Jc*_*ThT*_ and *Jm*_*ThT*_ and by *Jc*_*TMRM*_ and *Jm*_*TMRM*_. As an example, the flux equations is given here for ThT:

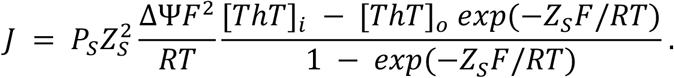

where, the indices *i* and *o* refer to the mitochondria and cytosol (in the case of mitochondrial flux) and cytosol and the cell exterior (in the case of plasma flux). *P*_*S*_ and *Z*_*S*_ denote the permeability and charge of the modelled dye, while *F, R*, and *T* denote the Faraday constant, the gas constant, and the temperature.

Finally, the relation between mitochondrial bound ThT level and ΔΨm is modelled through a linear function:

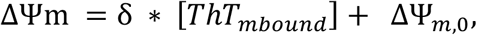

where *δ* is a scaling parameter and ΔΨ_*m*,0_ is the basal mitochondrial membrane potential.

The parameters used to simulate the system are provided in the MATLAB files given as *Supplementary File 1* and *2*.

## Acknowledgements

We would like to thank Dr. Ian Hands-Portman for his microscopy training and technical assistance.

## SUPPLEMENTARY FIGURES

**Supplementary Figure 1:**
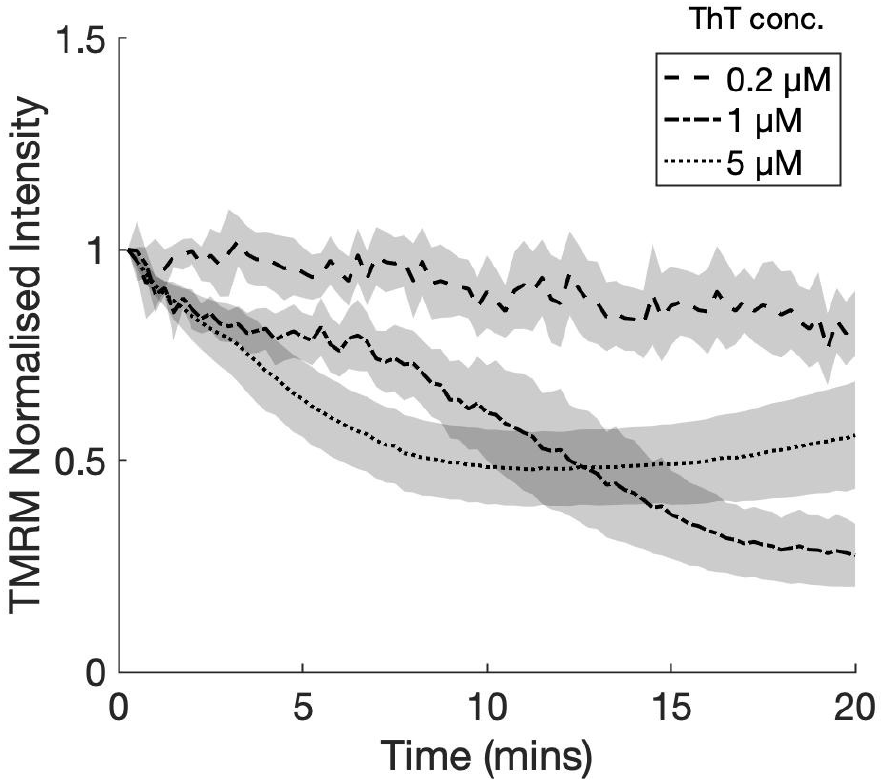
Shows the raw data from Fig 3B without background subtraction.

**Supplementary Figure 2:**
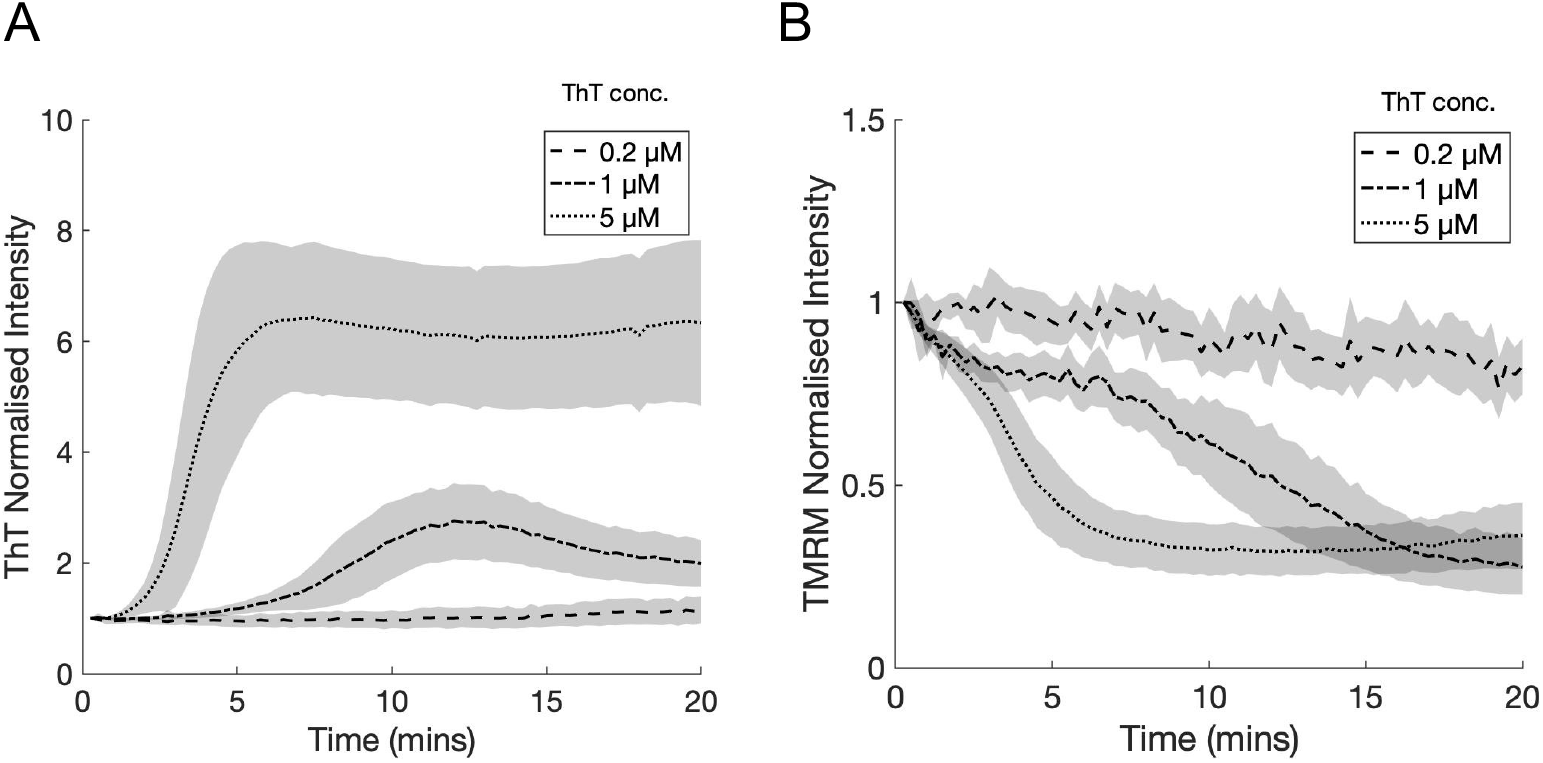
Nuclear ThT fluorescence **(A)** and cellular TMRM fluorescence**(B)** for cells co-stained with 25 nM TMRM and 0.2, 1 and 5 μM ThT and imaged in red and blue light. On each panel, the bold lines show the mean fluorescence intensity, and the shaded section shows population standard deviation, SD (mean±SD). Data is collated from three independent experiments, each with three technical repeats resulting in a total of 177 cells analysed.

**Supplementary Figure 3:**
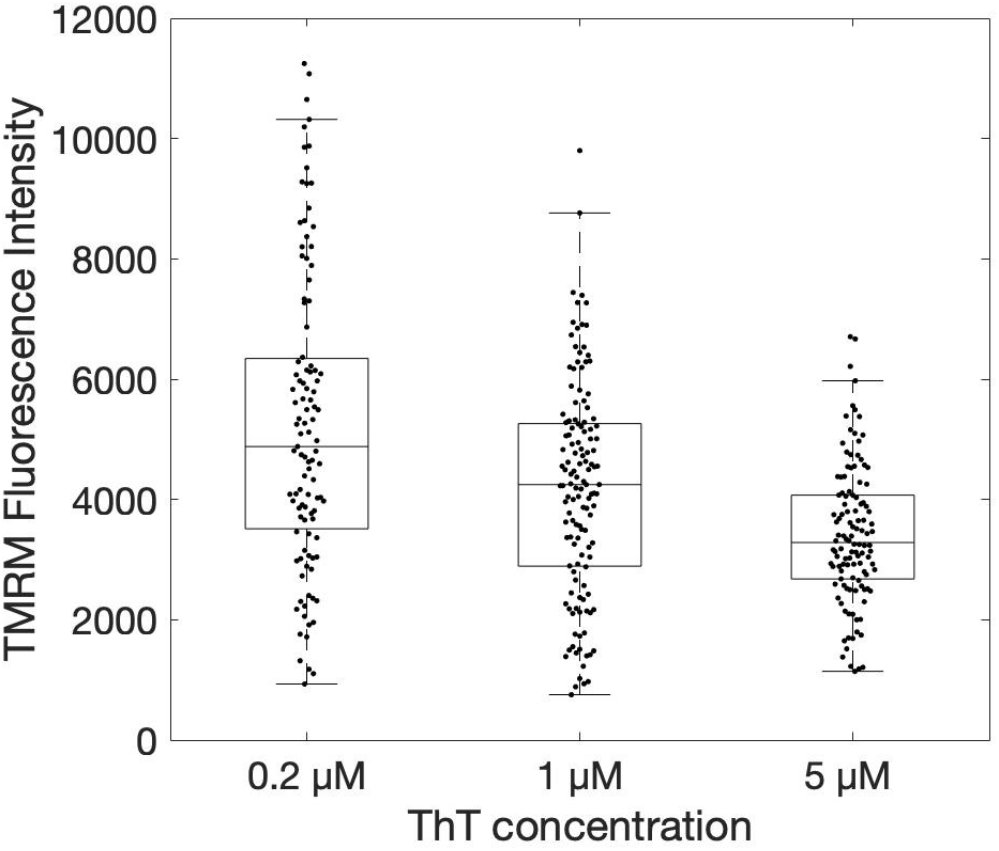
The response of cellular TMRM fluorescence (25 nM) after co-staining with 0.2, 1 and 5 μM ThT for 1 hr. Each data point represents one cell, with the analysed population for the three ThT concentrations totalling 107, 133, and 118 cells respectively.

**Supplementary Figure 4:**
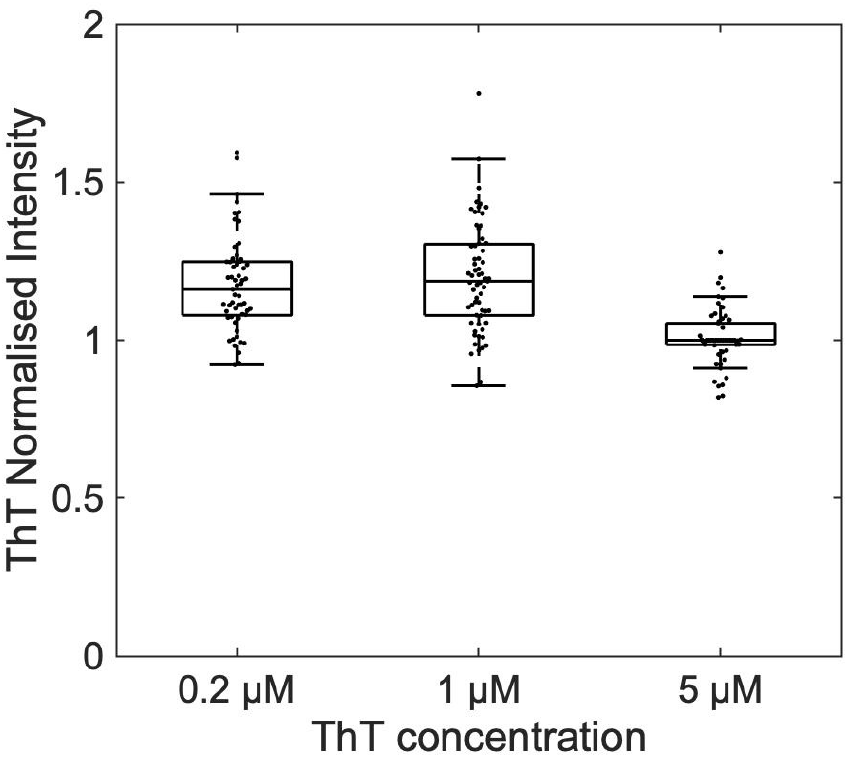
Cellular ThT fluorescence when cells were co-stained with TMRM (25 nM) and ThT (0.2, 1 and 5 μM). ThT fluorescence intensity ratio between before and after cells were imaged in red light for the time-lapse experiment shown in Fig 3C, i.e. beginning and end points of the experiment.

## Notes

### Competing Interest Statement

The authors have declared no competing interest.

